# Structural basis of phosphatidylserine enhancement of Gas6 ligand-induced Axl activation

**DOI:** 10.64898/2026.09.30.755801

**Authors:** Nicole Fournier, Camila Arango, Maria Lopez Vasquez, Pradeep Kumar Singh, Johnna Bingham, Huachen Gan, Adam W. Smith, Chrystal A. Starbird

**Affiliations:** Department of Biochemistry and Biophysics, University of North Carolina School of Medicine, Chapel Hill, NC, USA; Lineberger Comprehensive Cancer Center, University of North Carolina School of Medicine, Chapel Hill, NC, USA; Department of Chemistry and Biochemistry, Texas Tech University, Lubbock, TX, USA

**Author notes:** Co-first authors (Nicole Fournier, Camila Arango).

**Keywords:** Axl, Gas6, TAM receptor, cryo-electron microscopy, receptor tyrosine kinase, dimerization

## Abstract

TAMs (Tyro3, Axl, and MerTK) are receptor tyrosine kinases with primary roles in apoptotic clearance and immune suppression. In apoptosis, bridging complexes form between TAM receptors on phagocytic cells and TAM ligands bound to phosphatidylserine exposed on the surface of dying cells, but how these complexes form or why lipid is required for receptor activation is poorly understood. We present the first structures of a complete TAM receptor (Axl) and ligand (Gas6) 2:2 bridging complex with and without lipid, which suggest that the lipid functions to stabilize and promote active complexes. We also observe a stable 1:1 complex, presumed to be inactive, and provide evidence that these complexes are conversely able to form higher-order oligomers. We propose a mechanism for how lipids promote TAM receptor activation, providing a structural and biochemical basis for developing therapies that selectively target the lipid-dependent, immunosuppressive arm of TAM signaling exploited by tumors.

**GRAPHICAL ABSTRACT:** 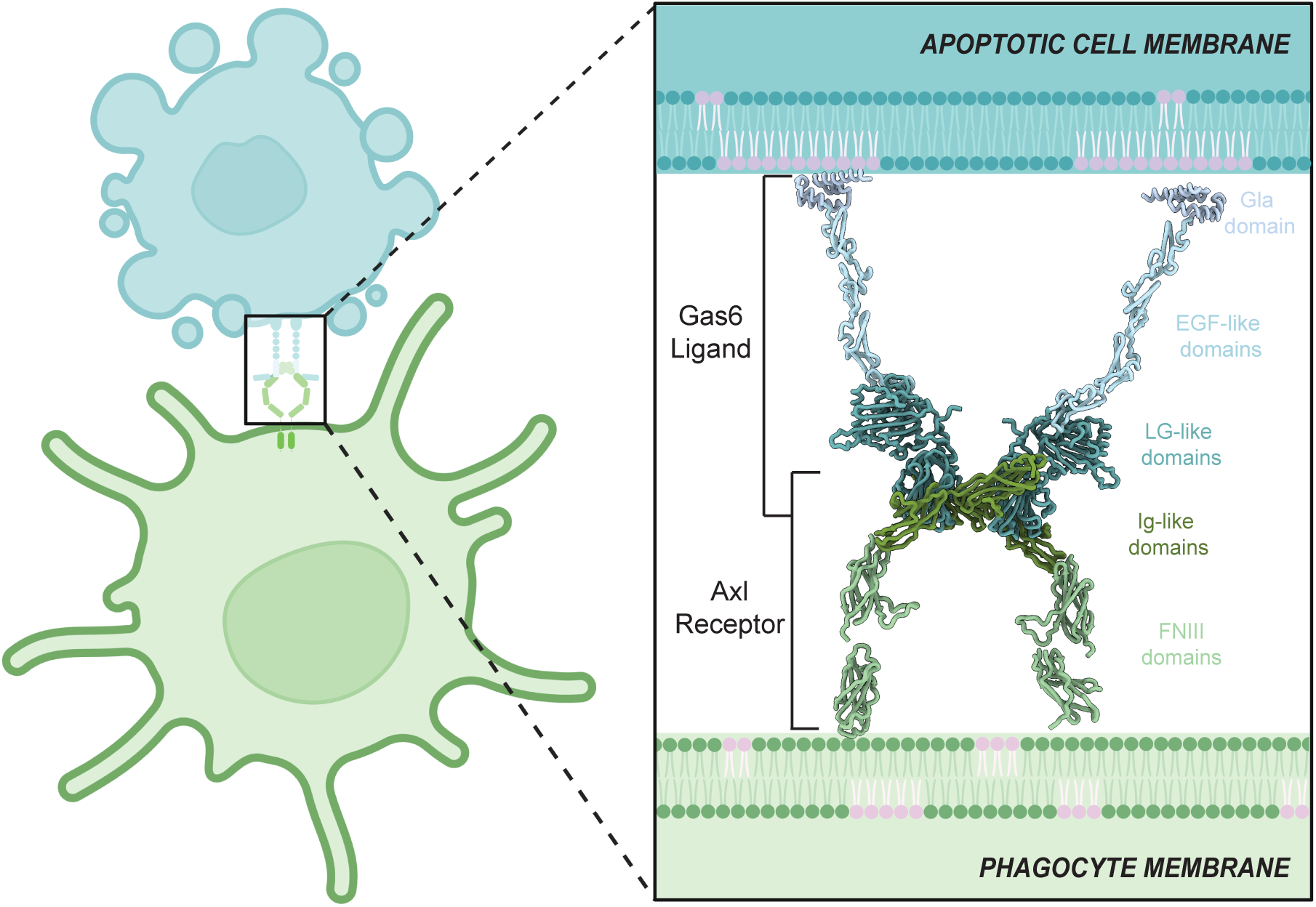

## INTRODUCTION

TAM receptor tyrosine kinases are key regulators of cellular homeostasis, mainly as drivers of efferocytosis involved in the phagocytic clearance of apoptotic cells by macrophages, dendritic cells, and other specialized phagocytes.^1^ TAM receptors also function as pleiotropic inhibitors of innate immune and inflammatory signaling triggered by Toll-like and type I interferon receptors^2,3^, they support platelet aggregation^3,4^, the maturation of natural killer cells^5,6^, and neuronal survival and myelination in the nervous system.^7,8^ Despite their critical roles in several regulatory processes and disease pathways, we know surprisingly little about the molecular details of TAM receptor:ligand structures and how TAM receptors interact with ligand to promote activation and subsequent signaling.

When cells are undergoing programmed cell death, the lipid phosphatidylserine is flipped from the inner to the outer leaflet of the plasma membrane and serves as an ‘eat me’ signal to phagocytes. This signal is recognized by the canonical TAM ligands, Gas6 and Protein S, which bind to exposed phosphatidylserine through residues γ-carboxylated in a vitamin-K dependent manner.^9^ TAM receptors can bind to ligand with or without lipid bound, and with affinities from the single digit nanomolar for Axl to hundreds of nanomolar for Tyro3 and MerTK. Axl activation, however, explicitly requires that ligand be bound to phosphatidylserine.^10,11^ Interestingly, while most receptor tyrosine kinase ligands exist as natural dimers that function to physically bridge two receptors, the TAM receptor ligands, Gas6 and Protein S, exist as monomers in solution.^12^ One possible hypothesis is that lipids facilitate dimerization of these ligands.

Though the specific, functional contributions of lipid binding are unclear, several diseases that involve TAMs also have known lipid contributions to disease pathology.^13-16^ In the tumor microenvironment, for example, cancer cells frequently display elevated levels of surface-exposed phosphatidylserine and secrete TAM ligands, essentially hijacking the apoptotic-clearance pathway to send an anti-inflammatory, immune silent signal even in the absence of true apoptosis.^17^ Downstream TAM activation in macrophages and dendritic cells induces the suppressor of cytokine signaling proteins SOCS1 and SOCS3, which shut down STAT1-dependent type I interferon and pro-inflammatory cytokine production^2^, while simultaneously skewing macrophages toward an M2-like, tumor-permissive phenotype marked by IL-10 and TGF-β secretion, impairing dendritic cell maturation and antigen presentation, and upregulating checkpoint ligands such as PD-L1.^18-22^ The net effect is a phosphatidylserine-driven signal to not ‘attack’, which dampens antitumor T-cell responses and contributes to resistance to immune checkpoint blockade — making the Gas6/Protein S–phosphatidylserine–TAM axis an active target for combination immunotherapy.^5,23^

Overall, a cell-to-cell bridging complex is thought to form in which ligand bound to exposed lipids from a dying cell interacts with the extracellular domain of a receptor expressed on the surface of a phagocyte.^24^ Graphical depictions of this complex are informed mainly by the dogma that receptor tyrosine kinases are activated by ligand-induced dimerization^25^, rather than direct knowledge of this assembly studied in TAMs. Here, we present the first structures that represent this entire cell-to-cell bridging complex, as well as studies that evaluate the specific role phosphatidylserine plays in structurally organizing this ligand:receptor assembly into an active complex.

## RESULTS

### Active complexes of Axl and Gas6 form in a concentration dependent manner

Although it’s not entirely clear what represents an active complex, a 2:2 complex of Axl and Gas6 is considered the minimal activating species, while a 1:1 complex (1:1 complex is defined here as a complex containing one copy of Axl and one copy of Gas6) is considered inactive.^11^ We completed mass photometry studies of purified full-length human ligand, Gas6, and Axl extracellular domain (ECD), which represent the domains associated with bridging interactions between cells. Overall, we observe that active (2:2) complexes of Axl and Gas6 form in a concentration dependent manner (Figure 1, teal distribution, Figure S1). At low concentrations (20-40 nanomolar), we observe either single species of Axl (52.4 kDa) and/or Gas6 (75.1 kDa), or average distribution peaks consistent with an inactive, 1:1 complex (127 kDa). As we move to higher concentrations, the population shifts entirely towards a 1:1 complex until we get to a concentration above 200 nanomolar, at which point there is a shift to a 2:2 (254 kDa) complex.

**Figure 1.**
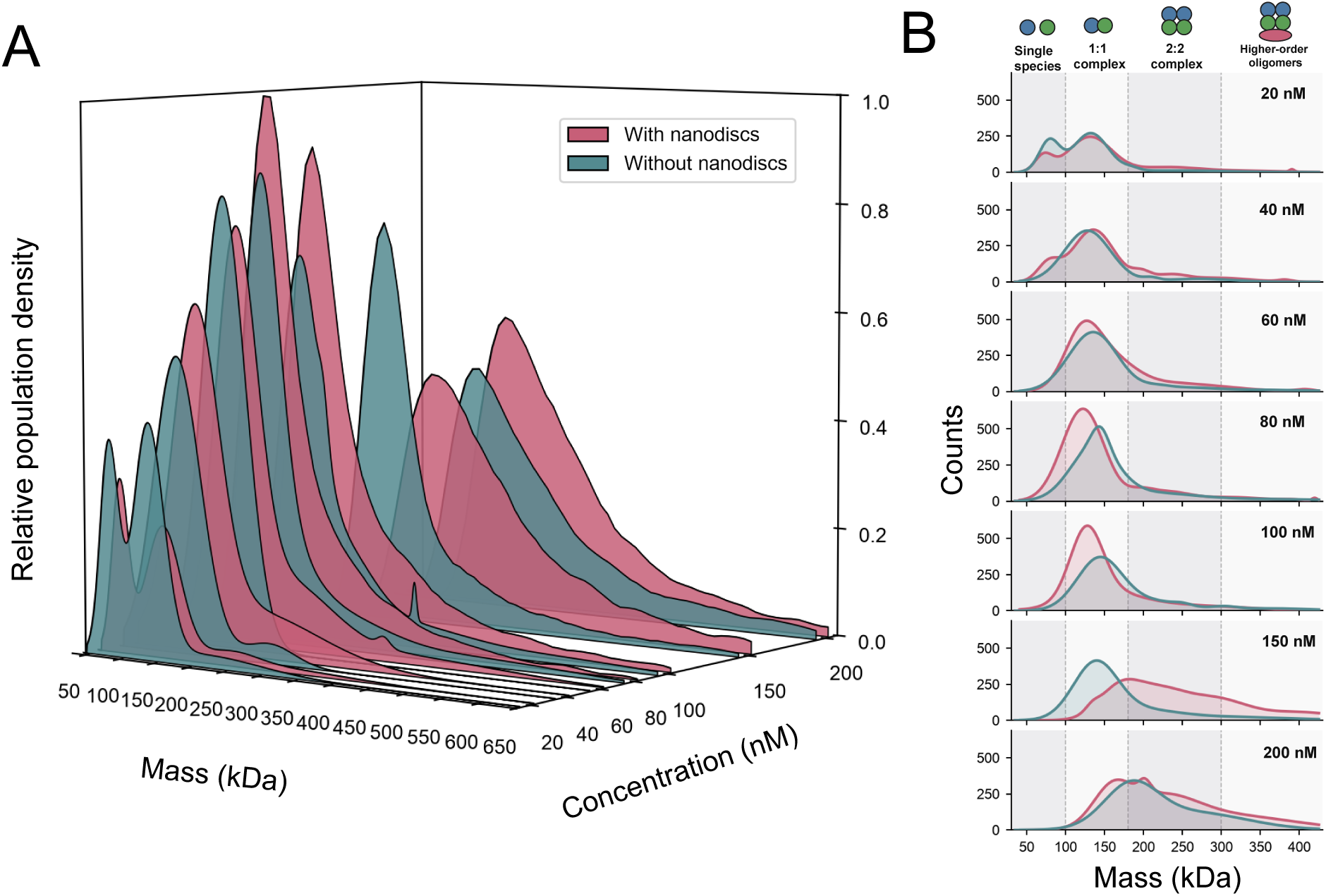
Mass Photometry reveals concentration-dependent assembly of higher order oligomers of Axl:Gas6 complex, enhanced by addition of nanodiscs containing 20% phosphatidylserine. **(A)** Three-dimensional ridgeline plot of mass photometry histograms for Axl:Gas6 mixed with nanodiscs (pink) versus without nanodiscs (teal) across increasing concentrations (20–200 nM). For each condition and concentration, histograms were created and averaged for three independent replicates, and a multi-Gaussian model was fit to the averaged distribution; curves are normalized so the global maximum across all conditions and concentrations equals 1, preserving relative differences in population size or counts. **(B)** Representative single-replicate mass histograms for the same concentration series, with mass regions corresponding to single species (30-100 kDa), 1:1 complex (100-180 kDa), 2:2 complex (180-300kDa), and higher-order oligomers (300-450 kDa) indicated above. Dots above represent Axl (green), Gas6 (blue) and nanodisc (pink oval). With nanodiscs present, higher-order complex formation increases at lower concentrations than for samples without nanodiscs.

Interestingly, when we add in nanodiscs containing 20% phosphatidylserine, we see an earlier shift, at ∼150 nanomolar, towards active complex and the presence of higher-order oligomers at relatively low abundance (Figure 1, pink distribution). While we do not see evidence of a stable nanodisc:ligand:receptor complex (which would result in a peak at ∼374 kDa, not observed), the presence of nanodiscs does appear to shift the population toward a 2:2 complex sooner than without phospholipids, which is consistent with our electron microscopy data (discussed later). Given its high affinity for Gas6 and the relatively high concentration of Gas6 in plasma^26^ and its broad expression in peripheral tissues, Axl has been proposed to remain constitutively bound to Gas6, with ligand binding alone being insufficient for receptor activation.^11, 27^ As such, this suggests that these inactive complexes are maintained on the cell surface in at low concentration and that lipids may function partially to help effectively concentrate the 1:1 complexes to form 2:2 complexes or higher order assemblies.

### Phosphatidylserine enhances Axl Activation

To determine how phosphatidylserine influences Axl activation in cells, we measured receptor and downstream signaling in response to Gas6 and PS-containing vesicles (Figure 2). We used NIH3T3 TKO cells, which do not express endogenous TAM receptors or ligands,^28^ that were engineered to stably express Axl (Figure S2). We then interrogate these cells with different activation protocols, including the addition of ligand and/or vesicles containing increasing amounts of phosphatidylserine (5-20%), and use phosphorylation of the receptor and downstream effector, ERK, as a readout for activation. We observe an increase in activation correlated to the addition of ligand, and a further increase as we add additional phosphatidylserine. This additional increase is not observed consistently downstream as phosphorylated ERK (two of three repeats show a similar increase), which may reflect a limitation of our experiment. For example, these experiments were halted at five-minute timepoints to optimize TAM phosphorylation signal based on investigative experiments in our lab, but this timepoint may not be ideal to observe significant changes in activation of downstream signaling. Additionally, although we cannot establish statistical significance due to the low sample number (n=3) and the variance from different westerns, the trend for each experiment is consistent and suggests that phosphatidylserine enhances Axl activation.

**Figure 2.**
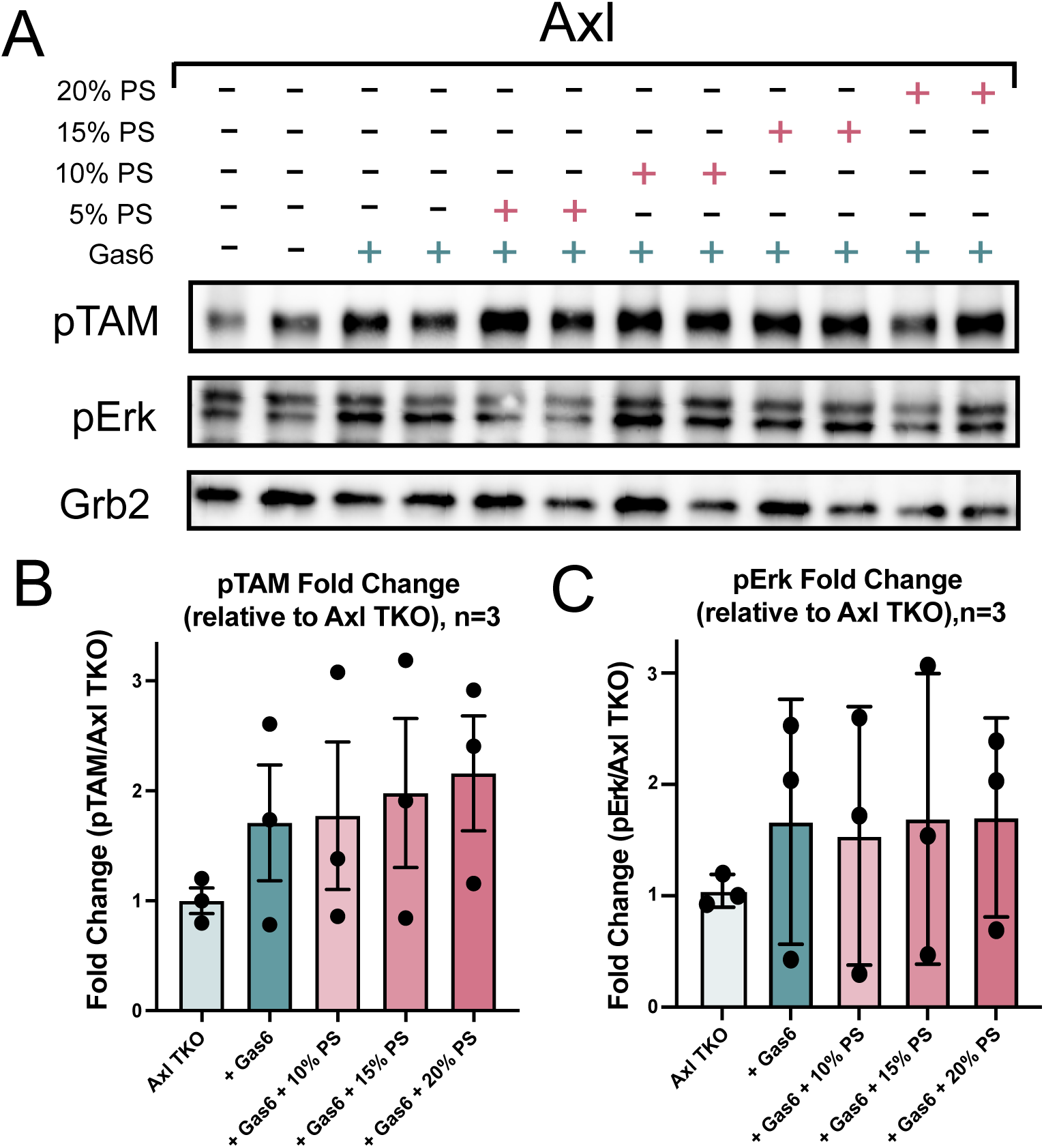
Phosphatidylserine enhances Axl activation in cells. (**A**) Representative Western Blot of cell lysates of NIH 3T3 TKO cells that stably express Axl with various conditions probing for phosphorylated TAM (pTAM) and downstream effectors, Erk. Grb2 serves as a loading control. Activation assay included several conditions represented by a blue cross to show addition of ligand and a pink cross the addition of vesicles. **(B**) Western blot quantification of a n=3 experiment, where fold change between band intensity of pTAM was divided against band intensity of Axl TKO alone to obtain fold difference between basal level of cells expressing Axl only to addition of ligand +/- vesicles. (**C**) Western blot quantification of pErk fold change (n=3), normalized to total Erk and calculated relative to basal Axl TKO cells, across the same ligand ± vesicle conditions shown in (**B**).

Phosphorylation levels for Axl with the addition of Gas6 alone are larger than for cells expressing Axl alone, and this is likely due to there being a basal level of cells expressing phosphatidylserine or undergoing cell death within these cultures. Overall, our data align with previous observations that phosphatidylserine is required for efficient Axl activation^10,11^ and support a model in which lipid engagement enhances signaling, at least in part, by stabilizing the active 2:2 Axl:Gas6 complex.

### Complexes of Axl and Gas6 are favored in 1:1 stoichiometry without phosphatidylserine

We determined three structures of an Axl:Gas6 complex without phosphatidylserine at a resolution range from 2.8-4 Angstroms (Figure 3A, S3, and Table S1). As depicted in the graphical abstract, Gas6 has seven total domains and Axl has four, making the entire predicted cell-to-cell bridging complex eleven domains. Gas6 is comprised of an N-terminal Gla domain, containing eleven predicted γ-carboxylated residues involved in binding phosphatidylserine, four epidermal growth factor (EGF)-like repeats, and two globular Laminin-G (LG)-like domains involved in interactions with receptor. The extracellular domain of Axl has two immunoglobulin (Ig)-like domains that are critical to its interaction with Gas6 and two membrane-proximal fibronectin type III (FNIII) domains. Overall, the structures we determined align quite well with the previously published minimal Axl:Gas6 crystal structure, which contains only the Ig-like domains of Axl and the LG-like domains of Gas6 (Figure 1B-D).^29^ We observe a 2:2 complex of Axl and Gas6 that includes density for three addition domains (1 EGF-like domain of Gas6 and the FNIII domains of Axl) not captured in the previous structure.

**Figure 3.**
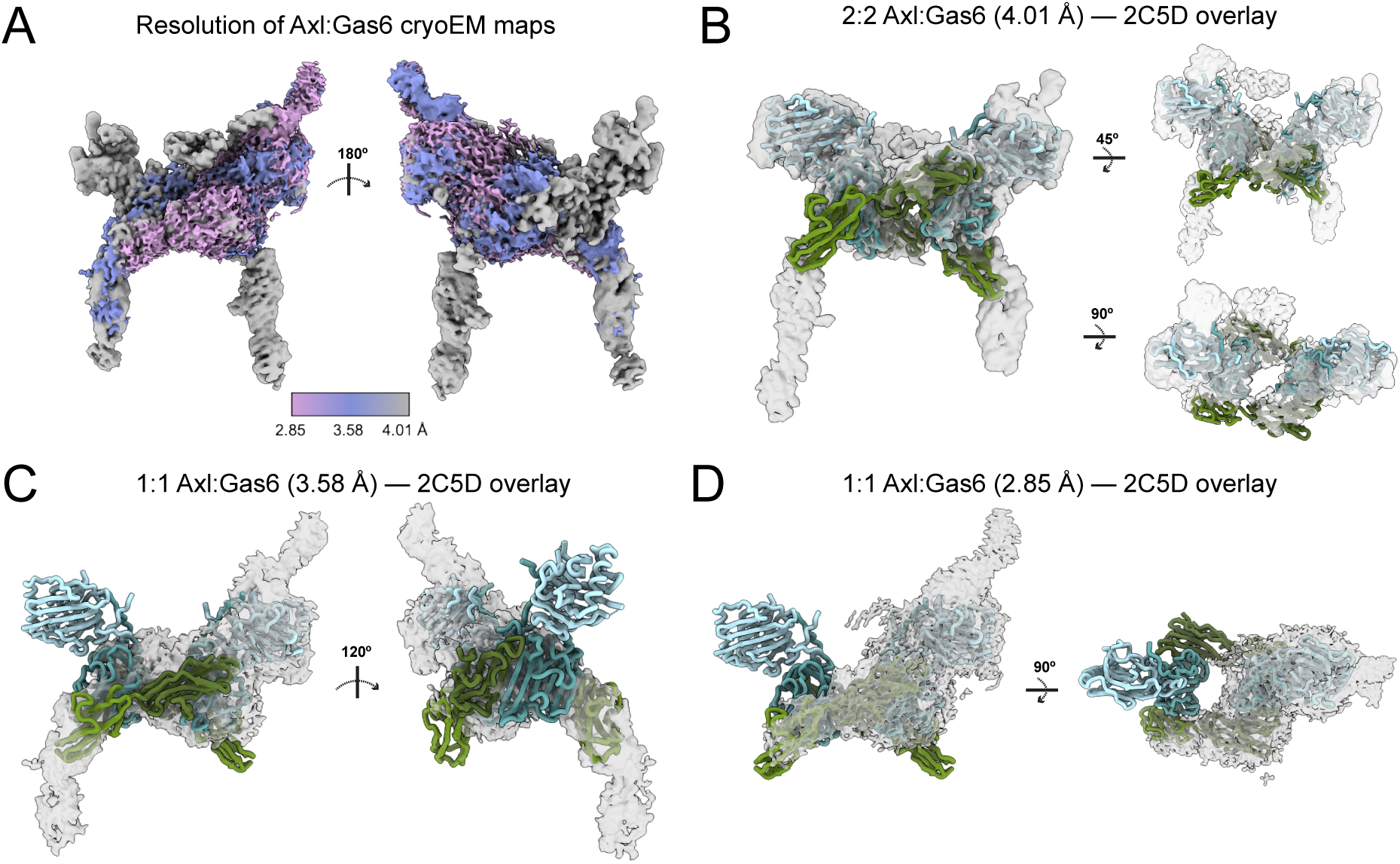
Cryo-EM structures reveal that Axl:Gas6 complexes favor 1:1 stoichiometry in the absence of phosphatidylserine. (**A**) Composite view of the cryo-EM densities for Axl: Gas6 structures without lipid, colored by local resolution (2.85–4.01 Å, scale bar) with 2.85 and 3.58 A being the monomer structures and 4.01 being the dimer structure. Composite view is shown at a second view rotated 180° about the y-axis. (**B–D**) Overlay of each determined structure with the previously published minimal Axl: Gas6 crystal structure (PDB 2C5D)^24^, shown at multiple rotational views as indicated. (**B**) 2:2 Axl: Gas6 complex (4.01 Å; contour level 0.00488), containing density for three additional domains not resolved in the minimal Axl:Gas6 crystal structure — one Gas6 EGF-like domain and both Axl FNIII domains — shown at three views. Additional density for three domains is segmented at a contour level of 0.025. (**C**) 1:1 Axl:Gas6 complex (3.58 Å; contour level 0.0891), shown at views related by 120°. (**D**) 1:1 Axl:Gas6 complex (2.85 Å; contour level 0.0793), shown at two views. In all overlays, Axl is shown in teal and Gas6 in green, with the corresponding cryo-EM density in transparent gray.

The majority of particles observed in these datasets (63% of 2D classes) are consistent with a 1:1 complex of Axl and Gas6, which is visualized in two of the structures we determined (Figures 3C-D). Both of these structures offer higher resolution views of two of the four EGF-like domains of Gas6 and the Axl FNIII domain adjacent the Ig domains. Interestingly, the 1:1 complex maintains the same overall orientation and fold observed in the 2:2 complex, creating an S-like bend in the overall structure (also observed in structures with lipid, discussed later).

Maintaining this orientation may be important to prime these complexes to readily oligomerize on the cell surface. Despite collecting a total of over seven million particles spread across datasets, we are only able to reliably visualize seven out of eleven total domains that represent the cell-to-cell bridging complex in these structures. This is likely due to the high level of flexibility in these complexes, which appears reduced in structures determined in the presence of phosphatidylserine membrane mimetics.

### Phosphatidylserine promotes the active 2:2 complex

In contrast to the more flexible complexes observed without lipid, phosphatidylserine-containing nanodiscs enabled the visualization of substantially more complete Axl:Gas6 assemblies. Cryo-EM revealed both 1:1 and 2:2 complexes containing density for all eleven domains of the predicted cell-to-cell bridging complex (Figure 4 and Table S1). We also obtained higher resolution structures of a 1:1 complex (3.5 Angstroms) and a dimer with unusual packing (at ∼3.4 Angstroms, Figure S4 and Table S1). For the lower resolution structures, AlphaFold^30,31^ was used to generate models of domains for which we did not have higher resolution information from our other datasets (namely the Gla-domain and 1 EGF-like domain of Gas6), and these were then placed into the maps.

**Figure 4.**
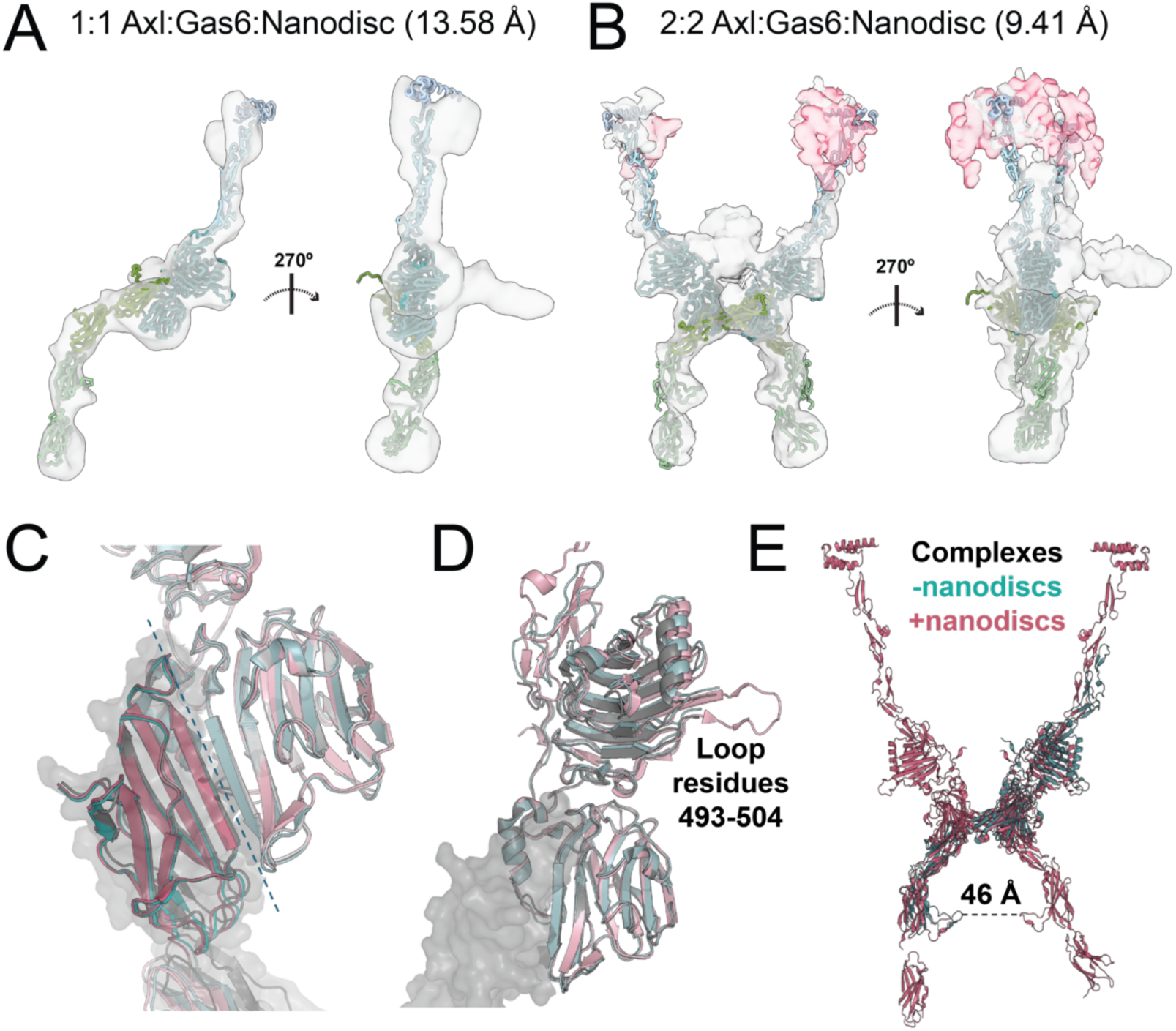
Cryo-EM reveals Axl:Gas6 complexes bound to phosphatidylserine-containing nanodiscs. **(A)** Low-resolution density of a 1:1 Axl:Gas6:Nanodisc complex (13.58 Å; contour level 0.187), with AlphaFold-predicted domains (Gla domain and one EGF-like domain of Gas6) fit into density not otherwise resolved, shown at a second view rotated 270° about the y-axis. An additional density is visible to the side of the complex, potentially indicative of clustering; no model is fit into this density. **(B)** Composite density of a 2:2 Axl:Gas6:Nanodisc complex (9.41 Å), shown at a second view rotated 270° about the y-axis, generated by segmenting and combining density from three separately contoured copies of the map: nanodisc density (pink; segmented using the Segger tool in ChimeraX, contour level 0.0134), EGF-like domain density (contour level 0.071, isolated using the Map Eraser tool shown in gray), and the remaining protein complex density (contour level 0.132, shown in gray). Two nanodiscs are resolved bound independently to the Gla domains of the two Gas6 arms. (**C**) Alignment of highest resolution structures with (pink) and without (teal) nanodiscs, aligned compared to 2C5D (grey). Dotted line splits the interface between LG1 of Gas6 and Ig1 of Axl. Grey surface outlines Ig1 and helps orient to the location of the Axl domain. (**D**) Similarly colored alignment of the same structures moved to highlight an LG2 domain loop not visualized in 2C5D or our structure without nanodiscs. Grey surface again orients to the location of the Axl Ig1 domain. (**E**) Visualization of model built for complex dimer with nanodiscs (pink) overlaid with high resolution monomer (teal) showing protrusion in FNIII domain of Axl.

In these lipid-bound structures, the population of the Axl:Gas6 complex shifts more towards 2:2 complexes when compared to the 2D classes of particles collected without lipids present. In datasets we collected with a higher concentration of nanodisc (∼1.4 nanodiscs:complex) we observe volumes for both a 1:1 complex in addition to a 2:2 complex, at higher frequency (66% of 2D classes). In both cases we observe density for all 11 domains representing a cell-to-cell bridging complex (Figure 4 and S5). The drastically reduced particle number required to observe stable 2:2 complexes that represent the entire bridging complex (<4000 particles compared to over 7 million) and the shift toward 2:2 complexes together support data from our mass photometry and cellular activation data to indicate that phosphatidylserine functions to stabilize active complexes. Importantly, for the structure that previews an active 2:2 complex, two nanodiscs are visualized bound separately to the Gla-domains from the two arms of the complex (Figure 4B). As such, the stabilization that allows us to visualize these domains is not due to being constrained by interaction with one nanodisc.

A careful analysis of all determined and modeled structures, including a comparison with the previous crystal structure, demonstrates that the most significant differences between these structures are indeed the interdomain angles, rather than the individual domain folds (Figure S6). When aligning the individual domains, the Ig1 domains across all structures has an average R.M.S.D. of 0.706 Angstroms, the Ig2 domains an average R.M.S.D. of 0.644 Angstroms, the LG1 domains an average R.M.S.D. of 0.477 Angstroms, and the LG2 domains 0.491 Angstroms. These values being well below 1.0 indicates that the majority of domains involved in the Axl:Gas6 interaction have highly similar folds. This is consistent with the conclusion that the impact of phosphatidylserine does not generally cause changes to the receptor and ligand core structures, but rather impacts the flexibility of the domains in a way that promotes formation of a stable, active complex.

When the highest resolution structure from data collected without (teal, PDB TBA) and with phosphatidylserine-containing nanodiscs (pink, PDB TBA) are compared, there are a few notable features that influence our thinking about the impact of lipids on these complexes. While the interface between Axl and Gas6 remains largely unchanged and overlays almost perfectly (Figure 4C), there is increased misalignment in the loops containing residues 104-113, as a result of a rotational shift in the domains. When we look further up the complex into the LG1 domain of Gas6, we see evidence for decreased flexibility in the structure with lipids, as several areas show increased secondary structure organization and density not seen in structures without lipids (or the previous crystal structure^29^) is able to be resolved (such as for loop residues 493-504, Figure 4D). Interestingly, we also observe a protrusion of primarily loop residues 264-283 in the least membrane-proximal FNIII domain (Figure 4E). This feature is observed in a high resolution 1:1 complex structure without lipid (PDB TBA) and in the model of the low-resolution volume of the complex with lipid (Figure 4B and 4E). A protrusion of this FNII domain is not seen in the recently published structure of the MerTK ECD or in models for the Tyro3 ECD^12^, making it likely that it’s a feature unique to Axl receptor. If we model the predicted distance between protrusions onto our dimer model, the distance is at least 46 Angstroms. An intriguing possibility is that this protrusion is involved in stabilizing active complexes, but this distance and our current structures do not support that hypothesis. It’s also possible that this feature is involved in interactions with alternative ligands or co-receptors not yet characterized.

### Phosphatidylserine promotes conformational changes that cluster FNIII domains to support activation

Key factors underlying activation of the intracellular kinases domains in receptor tyrosine kinases are the close proximity and reorientation required to support trans-autophosphorylation.^25,32^ In the ligand-induced dimer model and in expanded models of receptor clustering, kinase domain proximity is a driving factor for activation. For TAM receptors, given the overall structures presented here, bringing the kinase domains in proximity is likely to require structural changes that bring the FNIII domains closer, and is likely controlled by the hinge region between the receptor Ig domains. When we analyze our structures for those with the greatest predicted difference in FNIII domain range, the largest differences we observe are in the hinge region between the Ig domains.

Comparison of the hinge-region between Ig1 and Ig2 of Axl in structures with and without lipid show an average angular rotation in the lipid-bound structures (39 degrees in lipid-bound structures versus 22 degrees in lipid-free structures, Figure 5) in a direction that orients the receptor arm more towards the membrane and reduces the distance between the most membrane proximal FNIII domains. This conformational reorientation by phosphatidylserine is likely to reduce the distance between the intracellular kinase domains to facilitate trans-autophosphorylation, or activation of the kinases.

**Figure 5.**
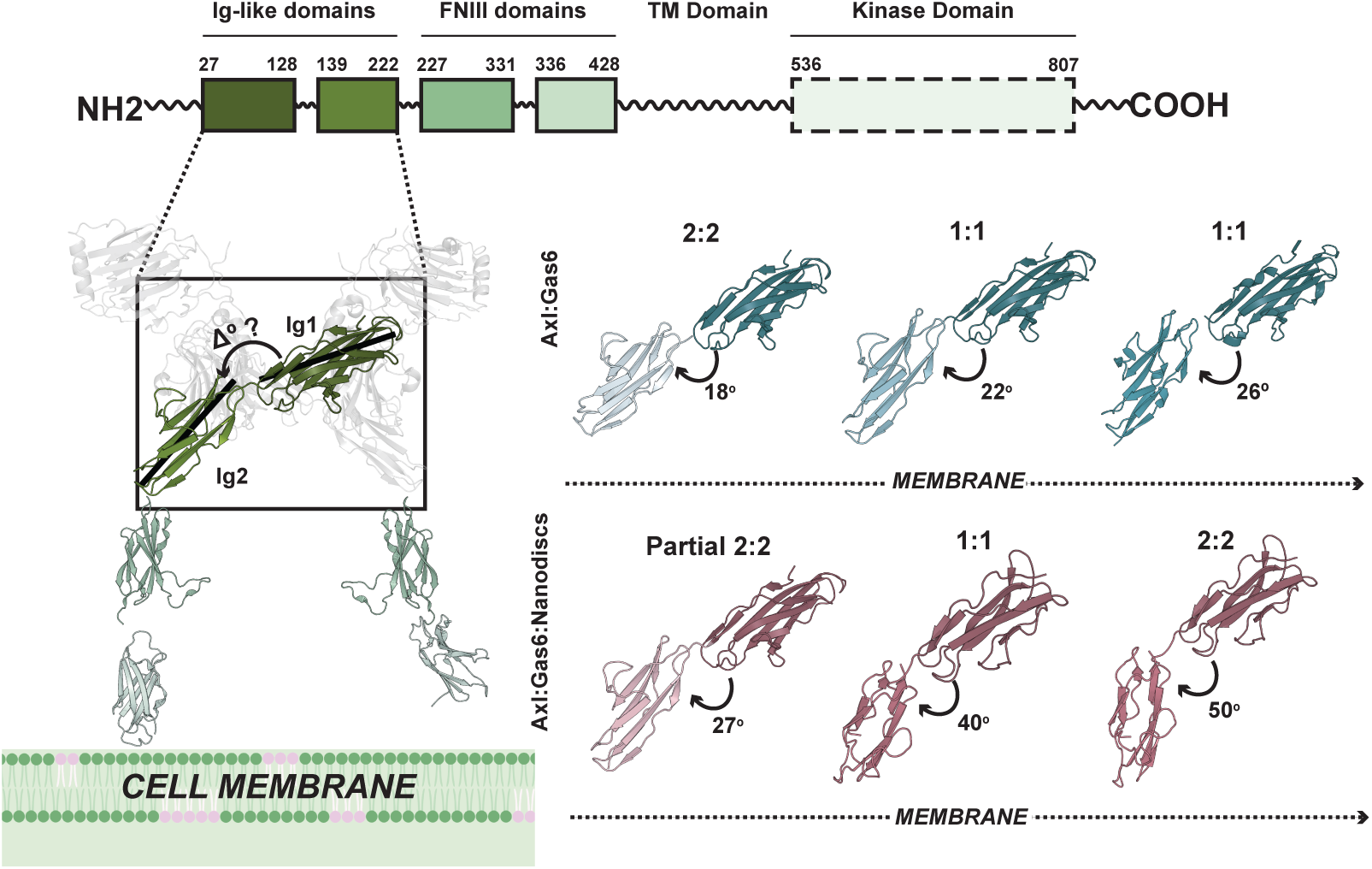
Phosphatidylserine-containing nanodiscs promote a hinge-angle reorientation between the Ig1 and Ig2 domains of Axl that orients the receptor toward the membrane. (Top) Domain architecture of Axl, indicating residue boundaries for the Ig-like domains (dark and light green), FNIII domains (teal and pale green), transmembrane (TM) domain, and kinase domain (dashed outline, not resolved in this study). (Bottom left) Structural model of the Axl:Gas6 complex with the Ig1-Ig2 hinge region boxed, illustrating the angle (Δ°) measured between the principal axes of the Ig1 and Ig2 domains. FNIII domains (green) are modeled using AlphaFold, as these membrane-proximal domains were not resolved at high resolution in contact with the membrane in our structures; dashed lines indicate the predicted trajectory of the Ig-like domains toward the cell membrane. (Bottom right) Comparison of Ig1-Ig2 hinge angles across individual Axl:Gas6 structures without lipid (teal, top row) and Axl:Gas6:Nanodisc structures with lipid (mauve, bottom row), arranged left to right in order of increasing (most negative to most positive) angular change relative to the reference structure (PDB 2C5D). In each structure, the darker shade (teal or mauve) indicates the Ig1 domain, positioned closer to the membrane, while the lighter shade indicates the Ig2 domain, positioned farther from the membrane. Angles are indicated below each structure, with the largest deviations reflecting the greatest conformational reorientation. Complexes bound to phosphatidylserine-containing nanodiscs show larger average hinge-angle rotations, in a direction that orients the Axl receptor arm toward the membrane, compared to structures without lipid. The principal (long) axis of each domain was calculated from Cα coordinates using singular value decomposition (SVD), a method for identifying the dominant direction of a set of points in 3D space, and the hinge angle was measured as the angle between the Ig1 and Ig2 axes. 2C5D was used as the reference/backbone structure for all hinge-angle comparisons.

### Gas6 promotes Axl assembly in live cells

We used PIE-FCCS to determine how Gas6 and phosphatidylserine (PS)-containing large unilamellar vesicles (LUVs) affect the organization and mobility of Axl in the plasma membrane of living cells (Figure 6).^33,34^ Measurements were collected across a broad range of receptor densities, from approximately 100 to 1,300 receptors/µm², with substantial overlap among all four experimental conditions. Mean receptor densities were 547, 412, 312, and 312 receptors/µm² for basal, PS-LUV, Gas6, and Gas6 + PS-LUV conditions, respectively.

**Figure 6.**
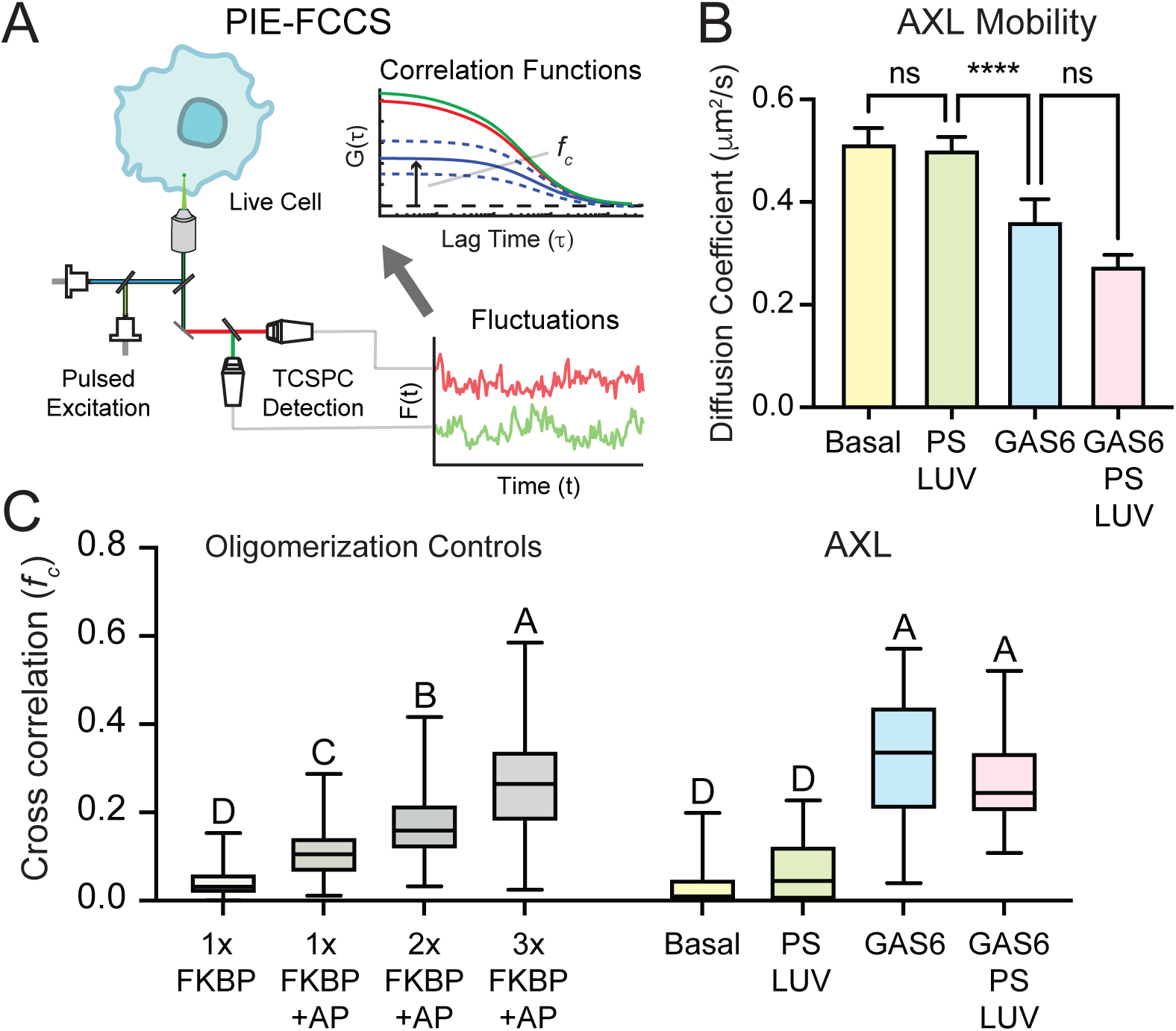
Gas6 promotes Axl oligomerization in live cells. **(A)** Schematic of PIE-FCCS. Fluorescence fluctuations from Axl-eGFP and Axl-mCherry fluorescent protein fusions were measured as receptors diffused through the observation volume at the plasma membrane. Auto- and cross-correlation functions were calculated from the fluorescence intensity traces, and the cross-correlation, *f_c_*, was used as a measure of receptor co-diffusion. **(B)** Axl diffusion coefficients measured under basal conditions or following treatment with PS-containing large unilamellar vesicles (PS LUV), Gas6, or Gas6 + PS LUV. Bars show mean ± SEM. Statistical significance was determined by Kruskal–Wallis test followed by Dunn’s multiple-comparisons test; ****P<0.0001; ns, not significant. **(C)** Axl *f_c_* values compared with previously characterized membrane-anchored FKBP oligomerization controls. Untreated 1×FKBP provides a monomeric control, whereas AP20187-treated 1×, 2×, and 3×FKBP constructs produce increasing levels of association. Axl exhibited low cross-correlation under basal and PS-LUV conditions, comparable to the monomeric control, whereas Gas6 increased *f_c_* to values comparable to the higher-order FKBP controls. Addition of PS LUVs did not further increase *f_c_* relative to Gas6 alone. Box plots show the median and interquartile range (25th–75th percentiles), with whiskers extending from the minimum to maximum values. Statistical significance was determined by Kruskal–Wallis test followed by Dunn’s multiple-comparisons test. Groups sharing a letter are not significantly different (P>0.05); groups with different letters are significantly different (P<0.05).

We examined receptor mobility using the diffusion coefficients obtained from the eGFP autocorrelation functions. Under basal conditions, Axl exhibited a mean diffusion coefficient of 0.51 ± 0.03 µm²/s (mean ± SEM), which was essentially unchanged following addition of PS-LUVs (0.50 ± 0.03 µm²/s). Gas6 stimulation reduced the diffusion coefficient to 0.36 ± 0.04 µm²/s, and Axl mobility was further reduced to 0.28 ± 0.02 µm²/s in the presence of Gas6 + PS-LUVs. Similar trends were observed for the mCherry-labeled receptor. Statistical analysis by Kruskal–Wallis test followed by Dunn’s multiple-comparisons test showed that Gas6 + PS-LUV treatment significantly reduced Axl mobility relative to basal and PS-LUV conditions, whereas PS-LUV treatment alone had no detectable effect (Figure 6B). The ligand-dependent decrease in mobility is consistent with formation of larger Axl assemblies following Gas6 stimulation, and potentially reflects the recruitment of downstream effectors following receptor activation.

We next quantified receptor co-diffusion using the fraction of correlation (*f_c_*) and compared the Axl measurements with previously characterized membrane-anchored FKBP oligomerization controls.^35^ Basal Axl exhibited little cross-correlation (median *f_c_* = 0.010), consistent with the monomeric FKBP control. PS-LUVs alone produced only a modest increase (median *f_c_* =.045) and remained statistically indistinguishable from the monomer control. In contrast, Gas6 caused a pronounced increase in Axl cross-correlation, with median (*f_c_*) increasing to 0.34 (interquartile range, 0.21–0.44). The Gas6-stimulated distribution overlapped the higher-order FKBP oligomerization controls, indicating formation of multimeric Axl assemblies. Addition of PS-LUVs to Gas6 did not further increase receptor cross-correlation (median *f_c_* = 0.25), and the Gas6 and Gas6 + PS-LUV conditions were statistically indistinguishable (Figure 6C).

Together, these measurements show that Gas6 shifts the mobile Axl population from a predominantly monomer-like state toward higher-order receptor assemblies in cells. Although PS-containing LUVs further reduce the mobility of Gas6-stimulated Axl, they do not produce a corresponding increase in *f_c_*. Thus, PS does not appear to increase the average oligomeric state of the mobile Axl population detected by PIE-FCCS, suggesting that its enhancement of Axl activation occurs without formation of larger receptor assemblies. Considered in the context of our mass photometry and activation data, this suggests that on the cell surface, Gas6 can dimerize Axl but these complexes are likely to be less stable and therefore less likely to promote activation without the presence of lipid, as supported by reduced diffusion in the presence of lipid.

## DISCUSSION

### Phosphatidylserine stabilizes and promotes conformational changes to activate Axl:Gas6 complexes

The work presented here, combined with contributions from other groups, allows us to propose a model for how phosphatidylserine modulates TAM receptor and ligand complexes in signaling. Based on previous work by others in the field, we know that binding of ligand alone is insufficient to activate Axl, and that Axl activation requires Gas6 be bound to phosphatidylserine, while MerTK and Tyro3 are partially activated without lipid.^10,11^ Our studies provide direct structural evidence that phosphatidylserine functions to oligomerize and stabilize active complexes to sustain signal transduction, and promotes conformational changes in Axl:Gas6 complexes that are likely to bring the cytoplasmic kinase domains into closer proximity for activation. In our model, Axl bound to Gas6 on cells is likely to exist as both 1:1 and 2:2 complexes, impacted by local concentration on the cell surface (Figure 7). We hypothesize that at a concentration predicated to occur in most cells, a higher percent of the population may exist as a 1:1 complex, but importantly, the presence of 2:2 complexes alone will not lead to activation. Consistent with this model, our single-molecule-sensitive PIE-FCCS measurements of full-length Axl in living cells show that Gas6 is sufficient to drive substantial receptor assembly, whereas addition of phosphatidylserine does not further increase the average multimeric state of the mobile receptor population under the conditions examined. These observations do not exclude a concentration-dependent effect of phosphatidylserine on the 1:1- to-2:2 equilibrium at lower receptor densities or in other membrane contexts, as suggested by the mass photometry data. We therefore favor a model in which phosphatidylserine primarily stabilizes and conformationally organizes activation-competent Axl:Gas6 assemblies, while potentially also shifting the 1:1/2:2 equilibrium under specific concentration regimes.

**Figure 7.**
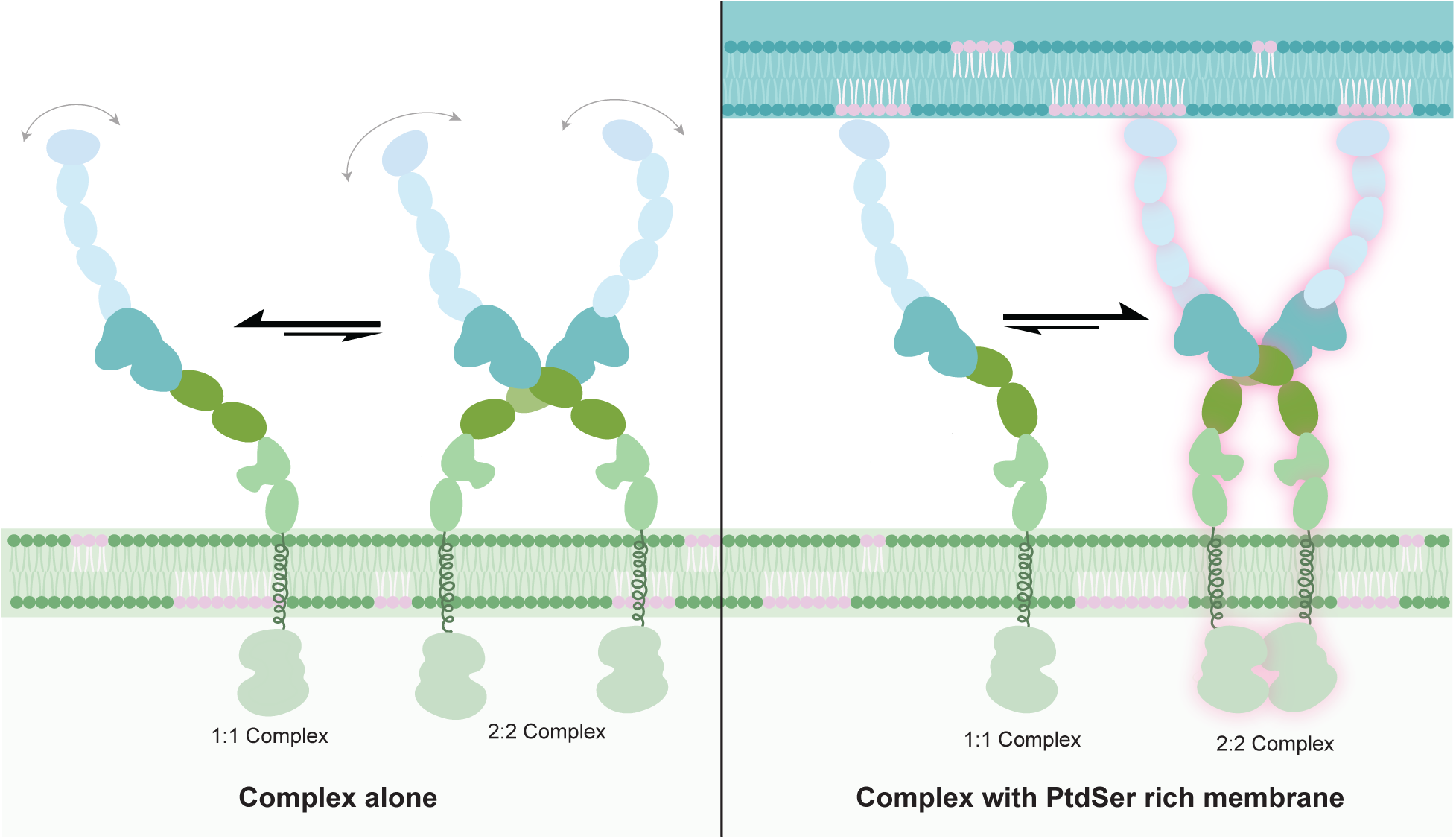
Proposed model for phosphatidylserine-dependent stabilization of the Axl:Gas6 bridging complex. Schematic depicting the equilibrium between 1:1 and 2:2 Axl:Gas6 complexes in the absence (left) or presence (right) of a phosphatidylserine-rich membrane environment. In the absence of a phosphatidylserine-containing membrane ("Complex alone"), the equilibrium favors the 1:1 complex over the 2:2 complex, with Gas6 (blue) unengaged and flexible. In the presence of a phosphatidylserine-rich membrane, Gla domain engagement with exposed lipid stabilizes the 2:2 complex (pink glow outline), representing the active, cell-to-cell bridging complex, and shifts the equilibrium to favor the 2:2 complex over the 1:1 complex. It is proposed that lipid binding both stabilize and tightens the complex, allowing the intracellular kinase domains to come into contact and become activated. Axl is shown in dark green (Ig-like domains) and light green (FNIII domains); Gas6 is shown in light blue (Gla and EGF-like domains) and teal (LG-like domains).

While both 1:1 and 2:2 complexes are likely to exist on cells where Axl is bound to Gas6, these complexes are likely to have a high degree of mobility and flexibility compared to those with lipid bound, precluding activation. As such, in our model, it’s only when these complexes are bound to phosphatidylserine from a nearby cell that we get a stable 2:2 complex where the FNIII domains shift closer together, due to a shift in the hinge rotation in the Ig domains towards the membrane, to promote trans-phosphorylation of the kinase domains. It’s less clear how phosphatidylserine might be dispersed in the lipid bilayer displaying an ‘eat-me’ signal and outside of the scope of this paper, but one could hypothesize that this model is more consistent with the idea that phospholipid forms clusters or high-density macro domains in phagocytic signaling.^36,37^

### Higher-order oligomer or cluster formation

The possibility that phosphatidylserine may form high-density regions on membranes corresponds well with the observation that TAM receptors are capable of forming higher order oligomers or clusters upon activation.^37^ The live-cell PIE-FCCS data support the formation of higher-order Axl assemblies following Gas6 stimulation, but do not show a further increase in average multimeric state upon addition of phosphatidylserine. We observe a low abundance of higher-order oligomers (larger than 2:2 complex) in our mass photometry data with and without nanodiscs, though these populations appear at lower concentrations in samples with nanodiscs.

Additionally, in of the cryo-EM structure data, we also observe density that we are unable to fully assign (seen in Figure 3B and Figure 4). This extra density is consistent with the start of another repeat of Gas6 and likely to represent a very low population of trimer or tetramer present in our datasets that we could not resolve. The idea that phosphatidylserine clusters support higher order arrays of receptor:ligand complexes remains speculative and although our data collectively do not exclude this idea, our findings more directly support a model in which higher-order assemblies can form with or without lipid, with phosphatidylserine potentially altering their stability or disruption at specific concentrations.

### Implications for MerTK and Tyro3 activation

Axl is the only TAM receptor that appears to have a strict requirement for phosphatidylserine engagement for activation by ligand: for MerTK and Tyro3 activation can occur without lipid present^11^, albeit at lower levels. We think the data herein would suggest, then, two potential scenarios for how lipid:ligand functions to activate these receptors. As ligand bound dimers can exist for all TAM receptors without lipid, in one scenario the ligand-bound complexes with Tyro3 and MerTK have increased stability compared to lipid-free structures of Axl and Gas6. In other words, lipid would not be an absolute requirement for stabilization of active complexes in these receptors.

In another scenario, lipid binding may not be required to promote the same conformational changes observed in our Axl:Gas6 complexes with lipid. One could imagine that the different orientations of MerTK and Tyro3 extracellular domains may be more permissive to self-activation, an idea supported by a recent paper from Starbird, et al.^12^ In this paper, it was observed that Tyro3 and MerTK appear to have interdomain angles that are inherently more curved towards the membrane, and this may make interaction with ligand:lipid assemblies less important to promote an active conformation. Either scenario causes us to wonder, then, how activation of MerTK and Tyro3 are controlled and leads us to hypothesize that one unifying possibility is that the surface concentration of both receptor and ligand-bound phosphatidylserine required for activation of the individual TAM receptors may be different. In this way, TAM receptors would not only be phosphatidylserine receptors, but finely tuned phosphatidylserine sensors. Regardless, it’s clear that TAM receptors have evolved to utilize lipid binding to phosphatidylserine for activation in manner that depends both on local receptor concentration and lipid-induced conformational changes.

## STUDY LIMITATIONS

Some of the volumes included in this study are very low resolution, particularly with electron microscopy data collected in samples that contain nanodiscs. However, given the highly similar domain folds (average R.M.S.D. of 0.615 Angstroms across all domains when individually aligned) in all high-resolution structure with and without nanodiscs, we believe the most critical differences between structures with and without nanodiscs is the interdomain angles, which are able to be interpreted utilizing these low-resolution volumes.

## ACKNOWLEDGEMENTS

We would like to thank members of the Earp Laboratory at the University of North Carolina School of Medicine for helpful discussions and feedback that aided in our studies. This research was supported directly by funding for our lab from the National Institutes of Health NIGMS [5R00GM144683], for the Smith lab from NIGMS [R35GM152126], and [T32-GM148376, for C.A. and M.L.V.]. Most of the cryo-electron microscopy work was performed at the Stanford-SLAC Cryo-EM Center (S²C²), supported by the National Institutes of Health Common Fund Transformative High Resolution Cryo-Electron Microscopy program under grant U24GM129541. We would specifically like to thank Grace Nye, staff at this facility who assisted greatly in electron microscopy data collection and early-stage processing. Part of this work was also performed in the UNC Protein Expression and Purification & Macromolecular Crystallography core lab supported by the National Cancer Institute of the National Institutes of Health under award number P30CA016086. Lastly, grid preparation and screening was completed at the National Institute of Environmental Health Sciences Cryo-electron Microscopy Core Facility, and the Cryo-Electron Microscopy Core at UNC Chapel Hill, supported by the National Cancer Institute of the National Institutes of Health under award number P30CA016086.

## AUTHOR CONTRIBUTIONS

N.F. and C.A. contributed equally to the article. N.F., C.A. and C.A.S. conceptualized the research. N.F., C.A., and M.L.V. expressed and purified proteins used in the study. C.A. and M.L.V. collected and processed mass photometry data. N.F and M.L.V. collected, and with C.A., J.B., and C.A.S., processed, analyzed, and validated all structural data. J.B. completed the activation assays and H.G. made cell lines used in these assays. A.S. and P.S. collected, analyzed, and interpreted all PIE-FCCS data and contributed explanatory text and figures corresponding to this technique. N.F. and C.A.S. wrote the paper, C.A. and C.A.S. designed figures, and all authors reviewed the paper and provided editing advice. This work was supported by funding acquired by C.A.S and A.S.

## METHODS

### Protein expression and purification

#### Gas6

Full-length human Gas6 was expressed from a pcDNA3.1(+) vector containing a C-terminal His^6^ tag and the native signal sequence. Recombinant Gas6 was expressed in Expi293 cells (Gibco, A14527) cultured in Expi293 expression media at 37°C with 5% CO_2_ in an incubator shaken at 125 rpm. Cells were transfected with a density of ∼3.0 x 10^6^ cells per milliliter using an Expifectamine 293 Transfection Kit (Gibco, A39251) in cultures supplemented with 10 µM Vitamin K (Ambeed, A763662). After incubation for 18-22 hours at 37C and 5% CO2 shaking at 125 rpm. Enhancers were mixed and added at recommended volumes for Expifectamine Transfection Kit. Cells incubated for a total of 84 hours before harvesting. Cells were removed by centrifugation and clarified conditioned media loaded onto Ni-Penta agarose column (Marvelgent Biosciences) using ∼1 ml of resin per Liter of culture. After washing with 0, 10, and 25 mM imidazole, protein was eluted with ∼150 mM imidazole in a buffer background containing 25 mM Tris-HCl, 150 mM NaCl pH 8.0. Fractions containing protein were pooled and diluted ∼6-fold in 25 mM Tris pH 8.0 to reduce the imidazole and salt concentrations to 25 mM or less for loading onto a 5 ml Source 15Q 4.6/100 PE column (Cytiva) for ion exchange.

Protein was eluted using a gradient of 25-250 mM NaCl/ 25 mM Tris at pH 8.0 over 5 column volumes. Desired fractions were pooled and concentrated using a 10 kDa MWCO Amicon Ultra Centrifugal filter before being loaded to a Superdex 75 size exclusion column for final purification. The Superdex 75 column was equilibrated using a buffer containing 25 mM Tris-HCl and 25 mM NaCl at pH 8.0. The molecular weight for Gas6 was estimated using sequence as well as information about the post-translational modifications, combined with measured molecular weights in both mass photometry and SAXS.^12^

#### Axl

The coding region for Axl extracellular domain was inserted into a pFastbacHT vector encoding residues 1-447 with the native signal sequence and a C-terminal His_6_ tag. For expression, 1-4L of Sf9 cells at a density of roughly 2 x 10^6^ cells/ml were transfected using recombinant baculovirus. Two days post transfection cells were harvested by centrifugation. Clarified conditioned media was loaded onto Ni-NTA Resin (Thermo) in a Kimble Konte Chromatography tube and equilibrated with 5 column volumes of 25 mM Tris-HCl, 150 mM NaCl pH 8.0 (buffer A). Following washes at 0, 10, and 25 mM imidazole in buffer A background, Axl was eluted using a stepwise method of buffer A containing 50, 75, 150 and 300 mM imidazole. Fractions were collected and samples analyzed via SDS-PAGE to confirm Axl protein location. Desired fractions were diluted 6-fold with 25mM Tris-HCL at pH 8.0 for a final concentration of 25mM NaCl and loaded onto a 5ml HiTrap Q HP column (Cytiva) for ion exchange. Protein was eluted using a gradient of 25-250 mM NaCl in 25 mM Tris at pH 8.0 for 20 column volumes. Desired fractions were collected and concentrated using a 10 kDa MWCO Amicon Ultra Centrifugal filter before being loaded to a Superdex 75 size exclusion column for final purification. The Superdex 75 column was equilibrated using a 25 mM NaCl, 25 mM Tris-HCl at pH 8.0. For PIE-FCCS, full-length AXL was subcloned into the eGFP-N1 and mCherry-N1 vectors using XhoI and AgeI restriction sites, generating constructs encoding AXL protein with C-terminal eGFP or mCherry tags (GenScript, Piscataway, NJ). The molecular weight for Axl was estimated using sequence, including the His_6_ tag, as well as information about the post-translational modifications, combined with measured molecular weights in both mass photometry and SAXS.^12^

### Preparation of MSP/Nanodiscs

MSP2N2 plasmid (Sigma MSP12) was transformed into BL21 competent cells and cultured in Lysogeny Broth (LB) in the presence of 50 ug/ml kanamycin. Induction with 0.5 mM IPTG occurred at an optical density between 0.6-0.8 at 37°C and a pellet was harvested by centrifugation. In Liter volumes, MSP2N2 purification involved thawing pellet on ice before adding 20 ml of lysis buffer (40 mM Tris-HCl, 300 mM NaCl pH 8.0, 1% TritonX-100 (w/v)) containing 50 units/ml of benzonase, 2 mg of lysozyme and 1 crushed protease inhibitor tablet (Sigma-Aldrich) and mixing until homogenous. Cells were lysed via sonication, and supernatant harvested by centrifugation and then applied to Ni-NTA resin equilibrated with 40 mM Tris-HCl, 300 mM NaCl pH 8.0, 1% Triton. Mixture was incubated with shaking for 1 hour at 4°C before being applied to a Kimble Konte Chromatography column set up for gravity flow. Nickel resin containing MSP protein was washed using 40 mM Tris-HCl, 500 mM NaCl, 1% Triton (wash buffer #1), followed by a wash with 40mM Tris-HCL, 500 mM NaCl, 50 mM Cholate, 5 mM imidazole pH 8.0 (wash buffer #2). The final wash step contained 40mM Tris-HCl, 500 mM NaCl, 10 mM imidazole pH 8.0 (wash buffer #3) before the protein was eluted in 40 mM Tris-HCl, 500 mM NaCl, 500 mM imidazole pH 8.0 and pooled. All wash steps were at least 10 column volumes. The desired protein was then dialyzed overnight into 20 mM HEPES, 100 mM NaCl pH 7.4. Samples containing MSP2N2 protein were then collected and concentrated to 18mg/ml to serve as MSP stock for nanodisc assembly.

Nanodisc assembly included measuring 2 µmol of DOPC lipids (Avanti) and 0.5 µmol DOPS (Avanti) that were dried out using nitrogen gas and left in vacuum desiccator overnight. Buffer (375 µL 11.25 mM Sodium Cholate, 20 mM HEPES, 300 mM NaCl pH 7.4) was added to dried lipids followed by a 10 second vortex and 5 min heat shock in 37°C water bath. This vortex/heating cycle is repeated 6 times before the lipid mixture is placed in a water bath sonicator for an hour. 125 µl of the MSP2N2 protein (at 18 mg/ml) is added to suspended lipid mixture and left to incubate for 20 minutes before being combined with 0.5 mg of Bio-Beads SM-2 Resin (BioRad) equilibrated in 20 mM HEPES, 300 mM NaCl pH 7.4. Lipid/MSP mixture is left to incubate with beads for 16-48 hours at 4°C. Nanodiscs were then purified using a Superose 6 size exclusion column equilibrated with 20 mM HEPES, 100 mM NaCl pH 7.4.

### Activation Assay

#### Cell Line Generation

Mouse Axl coding sequence fused with V5-tagged APEX construct was inserted into a 3rd-generation lentiviral transfer vector (adapted from Addgene Plasmid #193253) through NEBuilder HiFi DNA Assembly (New England Biolabs, #5520). To enable subsequent selection in cells, an IRES-puromycin-T2A-eGFP cassette was cloned downstream of the above coding sequence. Lentiviral pseudoparticles were packaged into Lenti-X 293T cells (Takara Bio, 632180) using packaging plasmids psPAX2 and pMD2.G, following the manufacturer’s protocol. The 48 and 72 h supernatant was filtered through a 0.2 µm PES filter (Thermo Scientific Nalgene, 725-2520) and used for the infection of NIH 3T3 TKO cells (obtained courtesy of the Nagata lab)^28^ in the presence of 8 µg/mL polybrene. 1 µg/mL puromycin (Corning, 61-385-RA) was added to the medium two days post-infection and maintained until positive colonies formed. The expression of Axl was confirmed by Western blot using Axl antibody (Cell Signaling Technology, #18554).

#### Liposome Preparation

For activation assays and PIE-FCCS studies, 10 mM vesicles were prepared containing 5%, 10%, 15%, and/or 20% DOPS in a DOPC background. Lipids were acquired from Avanti Research 18:1 PS (DOPS) CAS: 90693-88-2, LOT: 840035c-10MG-A-382, 18:1(Δ9-Cis) PC (DOPC), CAS:4235-95-4, LOT:850375C-25MG-A-342. Lipids were combined in a vial and dried with nitrogen gas and high vacuum overnight under light protection. Lipids were then resuspended in 10 mM HEPES, 150 mM NaCl pH 7.6 buffer. Resuspended lipids were frozen/thawed/sonicated/vortexed for at least 10 cycles. Lipids were then extruded with a PC Membrane 0.1 µm filter from Avanti Polar Lipids (SKU: 610005-1Ea, LOT: 150523Q2), 10 mm filter supports (SKU:610014-1Ea, LOT:151423Q2), using gas-tight syringes, 1000 µl (SKU:610017-1Ea, LOT: 151723Q2).

#### Cellular Assay

NIH3T3 TKOs stably expressing Axl were plated in a 6-well plate containing DMEM (Gibco, 11965092) product with 10% FBS (Gibco, REF-A52567-01 and Puromycin 1 µg/mL (Gibco, REF-A11138-03). Cells were serum-starved for 15 hours prior to the activation assay the next day. At 80-90 % confluency, 0.1 mL/well of activation medium was added to each well for 5 minutes (with or without Gas6 to a final concentration of 100 nM, PS vesicles 10 µM). After 5 minutes the activation mix was removed and the reaction stopped with the addition of cold lysis buffer (1x, Cell Signaling, 9803S) containing protease inhibitors and PhosStop at 1 tablet per 10 milliliters of lysis buffer solution (Sigma Aldrich, REF # 4906845001). Cells were harvested by scraping, vortexed for 30 seconds, stored on ice for 30 minutes, and vortexed again. Cell debris was removed by centrifugation and lysate was added to loading dye for gel electrophoresis.

Western blot: Samples were then run through a Bio-Rad 4-15% Mini-PROTEAN Stain-Free gel (4568086) for 60 minutes at 100 volts. Wet transfer to PVDF membrane was performed using Bio-Rad Mini Trans-Blot Electrophoretic Transfer cell. PVDF membrane was blocked for 1 hour in 5% milk/TBS-T. After washing in TBS-T the membrane was cut into thirds and blotted for pTAM (Cell Signaling 44463S), pERK1/2 (Cell Signaling 9106S), and Grb2 (Cell Signaling 3972) as a loading control. Secondary antibodies used (Anti-Rabbit IgG, HRP-linked Antibody from Cell Signaling # 7074S, Anti-Mouse IgG HRP-linked Whole Ab Cytiva # NA931V). Bio-Rad ECL (1705061) was applied to the membrane and incubated for 5 minutes and imaged using Bio-Rad ChemiDoc Touch Imaging system (Catalogue number #17001402J2P). Band intensity was quantified with Image Lab software.

### Pulsed interleaved excitation fluorescence cross-correlation spectroscopy (PIE-FCCS)

#### Cell Culture and Ligand Stimulation

COS-7 cells were cultured in Dulbecco’s Modified Eagle Medium (DMEM, Corning) supplemented with 10% fetal bovine serum (Thermo Fisher Scientific) and 1% penicillin– streptomycin (Thermo Fisher Scientific) under standard mammalian cell culture conditions. Cells were seeded onto 35-mm glass-bottom dishes (MatTek Corporation) and grown to 70–80% confluence before transient transfection with Lipofectamine 2000 (Thermo Fisher Scientific). A total of 2.5 μg of DNA, comprising mCherry- and eGFP-tagged plasmids in a 1:1 ratio, was used for each transfection to achieve balanced expression of both fluorescent tags and a final receptor density of approximately 100–1500 receptors/μm² at the plasma membrane. Approximately 20 h after transfection, the culture medium was replaced with phenol red-free Opti-MEM I Reduced Serum Medium (Thermo Fisher Scientific) before data collection.

For each construct, PIE-FCCS measurements were collected under both ligand-free and ligand-stimulated conditions using Gas6 (MedChemExpress) as the Axl ligand. Cells were first incubated with a range of Gas6 concentrations to identify the concentration that produced maximal, consistent receptor stimulation; a final concentration of 250 ng/mL was selected for all subsequent experiments, with or without co-addition of PS-containing large unilamellar vesicles (PS-LUVs). To stimulate receptor-expressing cells, a 500 μg/mL Gas6 stock solution was diluted to a working concentration of 250 ng/mL in Opti-MEM and added to cells approximately 15 min prior to the start of data collection; measurements were collected for up to 1 h following stimulation.

Control constructs designed to interpret fluorescence correlation (*f_c_*) values associated with monomeric, dimeric, and higher-order oligomeric states have been previously documented (2). These constructs were utilized to verify that the fluorescent protein tags do not inherently promote or inhibit dimerization. Based on these homotypic controls, *f_c_* values below 0.09 were interpreted as indicative of monomeric forms, those between 0.09 and 0.17 as dimeric, and values exceeding 0.17 as representative of higher-order oligomers (2).

#### PIE-FCCS Instrumentation, Data Collection, and Analysis

Single-color fluorescence intensity fluctuations were analyzed employing the time-time autocorrelation function; the correlation curves are presented on a semi-logarithmic time axis to comprehensively visualize the entire spectrum of relevant timescales within a single graphical representation. In summation, the amplitude of the autocorrelation function (ACF) exhibits an inverse relationship with the mean number of molecules present within the observation volume. In the context of two-color fluorescence cross-correlation spectroscopy (FCCS), two spectrally distinct fluorescent probes are activated and detected independently, thereby enabling each labeled population to produce its own ACF and allowing for the independent determination of the molecular density of each species. Specifically, in PIE-FCCS, the two excitation laser pulses are temporally phase-delayed relative to one another, facilitating the accurate attribution of each emitted photon to the corresponding excitation source and consequently eliminating spectral crosstalk among detection channels.

PIE-FCCS data were recorded utilizing pulsed interleaved excitation combined with time-correlated single-photon detection within a custom-designed inverted microscope. A pulsed supercontinuum laser (9.2 MHz repetition rate; SuperK, NKT Photonics, Birkerød, Denmark) was divided into 488 nm and 561 nm excitation beams through a series of dichroic filters and mirrors. To facilitate pulse interleaving, the two beams were routed via optical fibers of differing lengths, thereby introducing a specific delay between their arrival times at the sample. Prior to entering the microscope, the beams were recombined using a dichroic beamsplitter (LM01-503-25, Semrock) in conjunction with a custom filter assembly (zt488/561rpc, zet488/561m; Chroma Technology). The combined excitation beams were focused through a 100× TIRF objective onto a diffraction-limited spot at the peripheral plasma membrane of a COS-7 cell expressing the AXL-eGFP and AXL-mCherry constructs. Emitted photons were detected in a time-tagged, time-resolved mode by two avalanche photodiode detectors (Micro Photon Devices). System calibration, including the verification of confocal volume overlap between the two detection channels, was performed using a short, dual-labeled DNA standard prior to data acquisition. Additionally, positive and negative control samples were measured at the commencement of each session to verify the consistency of fit parameters.

Excitation beams were directed onto the flat, peripheral region of the plasma membrane to selectively assess membrane-bound receptors and to reduce the influence of fluorescence emissions from cytosolic vesicles or organelles. A single membrane area was analyzed per cell; each area was sampled six times consecutively, with each acquisition lasting 10 seconds.

Autocorrelation and cross-correlation curves were generated for each sample employing custom MATLAB scripts (MathWorks, Natick, MA). As previously documented (1, 4), a single-component, two-dimensional (2D) diffusion model was fitted to the average of the six consecutive 10-second acquisitions obtained for each membrane region.

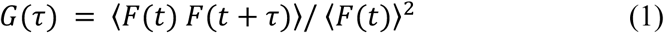

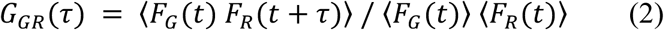

where *G*(τ) is the single-color autocorrelation function, *G_GR_*(τ) is the two-color cross-correlation function, and *δF*(t) represents the deviation of the fluorescence signal from its time-averaged mean at time *t*.

The intensity fluctuations were gated by the excitation pulse using PIE prior to calculation of both the autocorrelation and cross-correlation functions. In the autocorrelation analysis, the fluorescence intensity at time *t*, *F*(t), is compared with the intensity at a later lag time, *F*(t + τ); the resulting self-similarity, evaluated as a function of τ, provides quantitative information on particle number and diffusion. For each 10-s acquisition, intensity fluctuations were divided into 10-s bins and normalized to the square of the average intensity, consistent with Equation 1. The resulting autocorrelation curves were fit to a model describing two-dimensional diffusion in the membrane that incorporates a term accounting for triplet-state relaxation and other dark-state photophysics of the fluorophores (4):

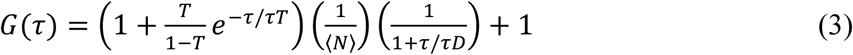

where *N* is the average number of diffusing molecules in the observation area, *τ_D_* is the characteristic diffusion time, *T* is the fraction of molecules occupying a dark or triplet state, and *τ_T_* is the corresponding relaxation time. The effective diffusion coefficient, *D_eff_*, was then calculated from the fitted diffusion time using the standard relation

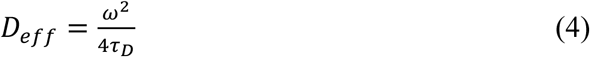

where *ω* is the radius of the effective observation area, determined from the DNA calibration measurements described below.

Finally, the fraction of cross-correlation, *f_c_*, a measure of the proportion of receptors that co-diffuse as a complex, was calculated from the amplitudes of the autocorrelation and cross-correlation functions of both fluorophores (2):

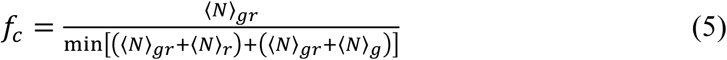

where *N_G_* and *N_R_* are the number of green-only and red-only diffusing species, respectively, and *N_GR_* is the number of co-diffusing (cross-correlated) species. Normalizing by the smaller of the two total populations ensures that *f_c_* ranges from 0 (no association) to 1 (complete association) regardless of any imbalance in the relative expression of the two fluorescent tags (2).

#### Control Samples for Fraction of Correlation (f_c_) Calculation

To calibrate *f_c_* values corresponding to defined oligomeric states, we used four membrane-anchored FKBP oligomerization control constructs prior to AXL data collection, as previously described (2). The untreated 1×FKBP construct served as a monomeric control, whereas treatment of the 1×, 2×, and 3×FKBP constructs with the small-molecule dimerizer AP20187 produced increasing degrees of association, corresponding to dimeric and higher-order oligomeric *f_c_* values, respectively (2). Laser powers for these control measurements were set to 0.350 μW (488 nm) and 0.80 μW (561 nm).

To calibrate the maximum achievable fluorescence signal and the effective confocal volume, we used a double-labeled DNA standard. A 40-nucleotide sequence with less than 50% G–C content (5′-ACA AGC TGG AGT ACA ACT ACA ACA GCC ACA ACG TCT ATA T-3′) was labeled with carboxy-tetramethylrhodamine (TAMRA) at the 5′ end and 6-carboxyfluorescein (FAM) at the 3′ end (Integrated DNA Technologies). The labeled strand was annealed with its complementary, unlabeled strand, added in molar excess, according to the manufacturer’s protocol, to generate the double-labeled DNA standard (TAMRA-40-FAM). This standard was diluted to a final concentration of 100 nM in 10 mM TE buffer. For calibration measurements performed in three-dimensional solution, laser powers were set to 7 μW (488 nm) and 7 μW (561 nm).

### Mass Photometry

Mass photometry measurements were performed using a Refeyn OneMP Mass Photometer (Refeyn Ltd, Oxford, UK) at a room temperature of 21° C. Microscope slides (High Precision Deckglaser) were cleaned through two rounds of sonication: 10 minutes in 50% isopropanol, and 15 minutes in Milli-Q water. After sonication, the beaker was rinsed and replaced with fresh Milli-Q water. An electric pump was used to dry the slides. Six-well reusable silicone gaskets (Grace Bio-Lab CultureWell gaskets, 50 diam x depth 3 mm x 1 mm) were placed on clean slides, and a drop of immersion oil (Olympus IMMOIL-F300CC for Microscopy) on the laser lenses was placed for visualization. Standards used for mass photometry measurements were Thermo NativeMark unstained protein standards (Thermo LC0725).

The Axl:Gas6 complex was measured with and without the presence of nanodiscs to determine the role of nanodiscs in higher-order complex stability. Axl, Gas6, and nanodiscs were individually run through ion-exchange chromatography and size-exclusion chromatography in buffer containing 25 mM Tris-HCl and 25 mM NaCl buffer at pH 8.0 (Buffer D). Peak elutions were pooled (Figure S1). Axl, Gas6, and nanodiscs were used at one-to-one molar ratios, and diluted corresponding to seven concentrations: in-excess of 200 nM and 150 nM; 100 nM; 80 nM; 60 nM; 40 nM; and 20 nM. Buffer D was used for droplet-dilutions, at a max volume of 20 µL. Movies of 5994 frames (59.9 second duration) were acquired for each sample, in triplicate using the Refeyn software (Refeyn Ltd, Oxford, UK).

Mass photometry data were processed and fit using a custom Python pipeline adapted from PyPhotoMol (version 1.0.1)^38^ (code available in the Supplemental/GitHub); the code was debugged and refined with the assistance of Claude (Anthropic). Single-particle contrast measurements were converted to mass using a calibration curve generated from NativeMark standards in DiscoverMP (Refeyn Ltd, Oxford, UK)For the 100, 150, and 200 nM samples without nanodiscs, whose exported files lacked calibrated mass values, masses were calculated from the contrast values using the linear calibration derived from the 20 and 40 nM measurements (mass = -20448.8245 × contrast - 0.257969864). For the averaged analysis, each of the three replicate measurements per condition and concentration was imported as calibrated single-particle mass events and binned into a mass histogram using identical bins (10 kDa bin width, 30-650 kDa window). The replicate histograms were then averaged bin by bin. Isolated single-bin artifacts, defined as a bin whose count exceeded five times that of both neighboring bins (consistent with unbound dye or optical noise rather than a true mass population), were clipped to the height of the adjacent bin. The averaged histogram at each concentration was fit with a multi-Gaussian model in PyPhotoMol. Peaks were detected automatically (minimum height of 5 counts, minimum separation of 4 bins, minimum prominence of 3 counts) and used as initial guesses, with fitted peak positions constrained to within 100 kDa and peak widths (σ) to within 150 kDa of the initial guesses. If a fit failed to converge, or collapsed into a degenerate spike (fitted maximum more than three times the maximum of the averaged histogram), it was repeated once with doubled tolerances. If it failed again, the averaged histogram was plotted directly and lightly smoothed with a Savitzky-Golay filter (window of 9 bins, third-order polynomial). This applied to the 80, 100, 150, and 200 nM samples with nanodiscs and the 150 and 200 nM samples without nanodiscs. For these curves, peak height and peak mass were taken from the smoothed averaged histogram (mass resolution limited to the 10 kDa bin width), and individual Gaussian component parameters are not reported. For all other curves, the position, σ, and amplitude of each fitted Gaussian, with their fit errors, were exported, along with the height and mass at the maximum of each curve. [State the fit-error threshold, e.g., "Only fitted components with position, σ, and amplitude errors of X% or less were interpreted."] To compare populations across concentrations and conditions, all curves were divided by a single global maximum (the tallest curve in the dataset, which was the smoothed averaged histogram for the 80 nM sample with nanodiscs), so relative differences in event counts are preserved. Curves were plotted in Python (Matplotlib) as a 3D ridgeline plot, with the two conditions offset slightly along the concentration axis so both remain visible.

To show the individual replicates without averaging (Figure 1B), each of the three replicates was analyzed separately. For each replicate, a separate figure was generated with one panel per concentration (20-200 nM), overlaying the with- and without-nanodisc fits. Each replicate was binned into a 10 kDa histogram (30-430 kDa window) and fit with a multi-Gaussian model using the same peak-detection, tolerance, and retry parameters as above. All 42 individual fits converged; 32 converged with the initial tolerances and 10 required the doubled tolerances, and no replicate required the smoothed-histogram fallback. The plotted curves are the fitted Gaussian sums for each replicate; the y-axis is in histogram counts per 10 kDa bin, with no averaging or density normalization, and is shared across panels within each figure. Despiking was not applied. Replicate numbers refer to the order of the three measurements at each concentration. Shaded mass regions (30-100, 100-180, 180-300, and 300-430 kDa) are visual guides marking the approximate ranges of single species, 1:1 complexes, 2:2 complexes, and higher-order assemblies [tie these boundaries to the expected masses of Axl, Gas6, and the nanodisc]. Final figures were assembled in Adobe Illustrator.

### Cryogenic Electron Microscopy

#### Vitrification

Following purification, samples were taken to the NIEHS Cryo-EM core facility for vitrification. Quantifoil Au 300 mesh R1.2/1.3 grids were plasma charged using an easiGlow (Pelco) at 0.50 mBar pressure with a glow time of 30 sec and a hold time of 10 sec. The Vitrobot Mark IV was set to 5 °C and 95-degree humidity. The purified protein complex at a concentration of 2-3 µM was pipetted into new, clean microcentrifuge tubes and spiked with 10% Fos-choline 8 (Anatrace) to help with orientation bias caused by the proteins interacting with the air-water interface. Grids containing nanodiscs came from two separate protein purifications. The first set of nanodisc/complex grids contained approximately 50 µl of 2 µM complex added to 50 µl of MSP2N2 nanodisc measured at 0.145 mg/ml. The second set of nanodisc/complex grids used 25 µl of MSP2N2 nanodisc measuring at 0.145 mg/ml and 75 µl of 2 µM complex. After vitrification, grids were transported in a CX100 transport dewar (Subangstrom) back to UNC Chapel Hill Cryo-EM Core for clipping. Copper washers and C-rings were provided by the core and used in the clipping process. Then grids were stored in long term storage dewars until screening.

### Data Processing and Structure Determination

All grids were screened at the UNC Chapel Hill CryoEM core before being sent to SLAC National Accelerator Laboratory for collection. Grids were collected in three different sets. All grids were collected on a Krios 300 kV microscope with a raw pixel size of 0.743, 2.7 mm spherical aberration and 50 e/A^2 total dose exposure. Movies were imported to CryoSPARC^39^ for processing. Each dataset was collected and processed independently from one another. After each collection on the Krios microscope, movies were imported to CryoSPARC and underwent standard Patch Motion Correction and CTF Estimation. The micrographs were then put through a Curate Exposure job to eliminate any problematic micrographs. Problematic micrographs in this case cover any micrographs with obvious alignment issues, broken spots or cracks, ice thickness issues, carbon edges, etc. During a manual curate job the CTF estimation values, relative ice thickness and motion distance were used as a baseline for curation. Any micrographs with ice thickness well beyond the mean value for that dataset were automatically excluded. Micrographs with a high degree of motion distance would present blurry and thus were excluded, and CTF values of higher than ∼10-12 were usually excluded. These parameters were adjusted based on each dataset.

After manual curation, micrographs were split into random subgroups (∼4-6 groups) using the exposure set tools to allow for more streamline processing and to distribute computation processing load across GPUs more efficiently. Each subgroup of micrographs was then run through a standard Micrograph Denoiser job to aid in visual inspection of particles during particle picking. To initiate particle picking a Blob Picker was used and set to 175-240 Angstrom size. Inspect Picks job was used interactively to identify “good” particles which were any particles that did not appear to be obvious contamination blobs. This job was followed by an Extract from Micrographs job which had varied box sizes between the lipid (640 box size) and non-lipid (384 box size) dataset to reflect the possible presence of lipid:ligand:receptor binding and ligand:receptor binding. An extensive and iterative round of 2D classes was performed in order to piece out different viewpoints of the 1:1 and 2:2 complexes. For more challenging views, a clean particle stack was curated and used in a Template Picker was used to reevaluate certain particle views. Once a clean particle stack was found, these particles would undergo *Ab-Initio* reconstruction to form an initial 3D map. From here, 3D maps were put through Homogenous or Heterogenous refinement (see Supplement Figures S3-S5 for which job refinement job performed on each structure) with C1 symmetry or C2 symmetry and relaxed symmetry pose marginalization to optimize the 3D model and filter out any junk particles that may be creating false positive signals. It should be noted that particle stacks for some structures remain very small due to extreme heterogeneity predominately seen in the distal domains. This low particle count would limit refinement abilities and resolution (see Figure S5). Lastly, a Non-Uniform Refinement job was performed on higher resolution structures to improve map quality for post-processing.

High resolution maps (3.5A <) were run through Model Angelo^40^ to get an initial reference model aligned to the map. This Model Angelo model was then brought into Coot and Isolde (UCSF ChimeraX) and compared against the only known partial structure of Axl:Gas6 (PDB:2C5D)^4^ and Alphafold predictions for Gas6 (Uniprot: Q14393) and Axl (Uniprot: P30530).^5^ For maps with obvious handiness issues, volumes were flipped and realigned to reference models and then built out using Coot and/or Isolde. For lower resolution structures (10A >), domain clusters were split using the predicted alignment error (pae_domain) and then docked into each low-resolution map using fitmap function in UCSF ChimeraX. Both low-resolution maps were flipped due to handiness before modeling. Domain clusters were set for Gas6 starting with the LG and EGF4 amino acid 239-677 followed by EGF3-1 amino acids 116-238 and amino acids 46-115 for the GLA domain. Domain clusters for Axl were set to IG domain with amino acids 26-224, followed by the first FNIII domain at amino acid 225-333 and amino acids 334-428 for the last FNIII domain cluster.

## KEY RESOURCES TABLE

(See STAR Methods Table)

## SUPPLEMENTAL INFORMATION

(separate PDF)

